# SMALL MOLECULE CLPP AGONISTS INDUCE SENESCENCE AND ALTER TRAIL-MEDIATED APOPTOTIC RESPONSE OF TRIPLE-NEGATIVE BREAST CANCER CELLS

**DOI:** 10.1101/2022.07.11.499620

**Authors:** Lucas J. Aponte-Collazo, Emily M. J. Fennell, Michael P. East, Thomas S. K. Gilbert, Paul R. Graves, Hani Ashamalla, Edwin J. Iwanowicz, Yoshimi Endo Greer, Stanley Lipkowitz, Lee M. Graves

## Abstract

Imipridones are a novel class of anticancer drugs with promising antiproliferative effects in several cancer cell types, including breast cancer. Recent studies identified the mitochondrial ATP-dependent caseinolytic peptidase P (ClpP) as the target for imipridones and related analogs. Despite these findings, the specific processes by which ClpP activators inhibit cancer cell growth remain poorly understood. Here we report that two structurally distinct ClpP activators, ONC201 and TR-57, promote senescence in SUM159 and MDA-MB-231 triple-negative breast cancer (TNBC) cell lines. Induction of senescence was measured through β-galactosidase assays and confirmed by the increase of H2A.X phosphorylation, hypophosphorylation of retinoblastoma protein (Rb), upregulation of multiple interleukin mRNAs and other markers. The level of senescence induced by these compounds was equivalent to that observed with the CDK4/6 inhibitor and positive control abemaciclib. To confirm the crucial role of ClpP activation in senescence induction, we generated ClpP null TNBC cell lines using CRISPR interference (CRISPRi). Neither ONC201 nor TR-57 induced senescence in the ClpP null models. Incubation of WT cells with ClpP activators led to a reduction in the levels of apoptosis-related proteins like XIAP, SMAC/DIABLO, Survivin, DR4 and DR5, which correlated with the lack of apoptosis observed in these cells. Interestingly, treatment with TR-57 strongly reduced apoptosis induced by staurosporine but increased sensitivity to tumor necrosis factor-related apoptosis-inducing ligand (TRAIL). To investigate the enhanced effects of TRAIL, we examined the expression of Wee1 in senescent cells and found that both TR-57 and abemaciclib down-regulated Wee1. Addition of a Wee1 inhibitor partially sensitized cells to TRAIL suggesting the importance of Wee1 in this process. In summary, we show that ClpP activators induce senescence in a ClpP-dependent manner and that combined treatment of ClpP activators with TRAIL provides an effective approach to eliminate malignant senescent cells *in vitro*.

**HIGHLIGHTS:**

- Treatment of TNBC cells with ClpP activators induces senescence in vitro
- Induction of senescence is ClpP dependent
- Activation of ClpP leads to changes in mRNA levels of senescence associated cytokines
- Senescent TNBC cells are sensitized to TRAIL mediated apoptosis

## Introduction

Triple-negative breast cancer (TNBC) is the most aggressive subtype of breast cancer with a significantly lower survival rate than other subtypes^1^. Since TNBC cells lack estrogen and progesterone hormone receptors or enrichment of the receptor tyrosine kinase HER2, targeted therapeutics like tamoxifen and trastuzumab are largely ineffective. Current treatments for TNBC patients are limited to surgical intervention and traditional chemotherapies accompanied by immunotherapies^2,3^. ONC201 is a recently discovered imipridone molecule in Phase I/II clinical trials for a large variety of aggressive cancers including breast, glioblastoma, and endometrial among others^4,5^. Since its discovery, multiple ONC201 analogs have been synthesized including ONC206, ONC212 and the highly potent TR compounds^6–10^. While ONC212 has shown promising broad-spectrum activity across multiple tumor types *in vitro*, studies to facilitate approval of first-in-human clinical trials are currently ongoing. Recently, ONC206 entered clinical trials for the treatment of central nervous system cancers^11^.

Although ONC201 was initially proposed to promote apoptosis through increased TRAIL formation and signaling^12^, its growth inhibitory effects were shown even in the absence of TRAIL in breast cancer cell lines^13^. The direct target for ONC201 and related compounds was unknown until our lab and others demonstrated that they bound and activated the mitochondrial ATP-dependent Clp protease (ClpP), the catalytic subunit of the ClpXP complex ^9,14^. ClpP proteolytical activity is regulated by ClpX in an ATP-dependent manner. Ishizawa et al. demonstrated that ONC201 displaced ClpX and converted the ClpP subunit to an open and active conformation. In addition to its well-established role in mitochondrial protein degradation, ClpP activation has been linked to regulation of cell growth and apoptosis through signaling events like the unfolded protein response and the integrated stress response^9,13,15–19^. Studies from our lab and others showed that while ClpP activators reduced TNBC cell proliferation, they did not significantly induce apoptosis *in vitro*^9,13^. Since growth arrest and lack of apoptosis are typically observed in senescent cells^20,21^, we investigated whether ClpP activators were inducing senescence in TNBC models.

Cellular senescence is characterized by a lack of cell cycle progression, increased levels of β-galactosidase (β-gal), increased levels of DNA damage, high levels of cyclin-dependent kinase inhibitor p16INK4a, elevated expression of anti-apoptotic proteins, and apoptosis resistance among others markers^22–24^. Additionally, senescent cells display activation of senescence-associated secretory phenotype (SASP) which is believed to enhance innate and adaptive immune cell recruitment by secretion of specific cytokines and chemokines^25,26^. Cell senescence *in vitro* can also be induced by mitochondrial dysfunction. Mitochondrial dysfunction-associated senescence (MiDAS) shares many similarities with other types of senescence but has a different SASP profile than other types of senescence^27,28^.

Induction of senescence as an alternative strategy to reduce tumor growth, has been the focus of multiple studies^29,30^. This has led to the approval of senescence-inducing drugs, like palbociclib and other CDK4/6 inhibitors, as a combinatorial treatment for specific cancer types including TNBC^31^. While therapy-induced senescence was initially thought to be an effective standalone treatment to inhibit cancer cell growth, further research has shown that chronic presence of senescent cells can be detrimental to tumor reduction since it promotes inflammation and modulates metastasis of nearby non-senescent cells^32^. Thus, these studies highlight the need to better understand the mechanisms promoting senescent cell clearance after therapy-induced senescence.

In this study we show that a recently identified ClpP activator, TR-57, induces senescence in a ClpP-dependent manner. We also demonstrate that TR-57 generates a moderate SASP profile and activates AMPK and other senescence-associated events similar to MiDAS. Lastly, we demonstrate that ClpP-activated senescent cells have lower levels of key apoptosis-related proteins and reduced apoptosis, but show increased TRAIL-mediated apoptosis. These results suggest that combining ClpP activators with TRAIL agonists may be an effective treatment approach for TNBC.

## Materials and Methods

### Chemicals

ONC201 was obtained from SelleckChem. The TR-57 and TR-107 compounds were supplied by Madera Therapeutics, LLC. abemaciclib was obtained from advanced Chemblock inc. TRAIL (Cat. No. HY-P7306) and Adavosertib (Cat. No. HY-10993) were obtained from MedChemExpress.

### Cell Culture

Human TNBC cell line SUM159 were a generous gift from Dr. Gary Johnson at UNC CH. MDA-MB-231 were a generous gift from Yoshimi Greer at NCI. SUM159 cells were cultured in Dulbecco’s modified Eagle’s medium: Nutrient Mixture F-12 supplemented with 5% fetal bovine serum 5 μg/mL insulin, 1 μg/ mL hydrocortisone, and 1% mixture of antibiotic-antimycotic. MDA-MB-231 cells were cultured in RPMI 1640 media supplemented with 10% FBS and 1% antibiotic–antimycotic.

### Senescence-associated β-galactosidase staining

SUM159 and MDA-MB-231 cells were stained for SA-β-gal detection using a Senescence β-galactosidase Staining Kit (#9860, Cell Signaling). Staining was performed by following the manufacturer’s instructions and incubating the stained samples for 48hrs at 37°C without CO2. Images for SA-β-gal quantifications were acquired using a ZEISS Axio Vert.A1 inverted microscope at 10x magnification. For each biological replicate, an area of the well was randomly selected, and the total of SA-β-gal positive cells were manually counted. Quantification and further analysis of the data was performed using Fiji software (ImageJ).

### Viability assays

#### MTS

Cell viability assays were performed by plating SUM159 WT or SUM159 ClpP null cells (1000 cells/well) on a 96-well plate (655-180, Greiner) in their respective media as indicated previously. Cells were allowed to adhere overnight. After adherence, the media in each well was aspirated and replaced with 100 μL of media containing the indicated compound(s). Cells treated with DMSO (vehicle) were used as a negative control in all experiments. Cells treated with the indicated concentrations of selected compounds for 72 h in 100 μL of incubation media were supplemented with 20 μL of 0.6 mM resazurin (Acros Organics 62758-13-8) and incubated for 30 min at 37°C. 75 μL of each sample was then transferred to a black 96-well plate (CLS3915, Millipore Sigma), and the relative fluorescence of resorufin across samples was determined using a PHERAstar plate reader (BMG Labtech) with fluorescent module FI: 540-20, 590-20. The results were analyzed using GraphPad Prism 9 software.

#### Total Cell count

Total cell counting assays were performed by plating and treating cells as described above. At the predetermined time points (0, 24, 48, or 72 h), media was aspirated, and 100 μL of Hoechst stain (1 μg/mL, H3570, Thermo Fisher Scientific) was added to each well and allowed to incubate for 15 min at 37°C. Total cell number was then quantified using the Celigo Imaging Cytometer (Nexcelom).

### Immunoblotting

Cells were plated in a 6 well plate (100,000 cells/well) or 10 cm dishes (1,000,000 cells/plate) and treated with compounds as described above. Following treatment, cells were lysed with RIPA buffer [no SDS, 2 mM Na(VO3)4, 10 mM NaF, 0.0125 μM calyculin A, and complete protease inhibitor cocktail (11873580001, Roche Diagnostics)] and lysates immunoblotted as described previously^9^. Nitrocellulose membranes were incubated with the indicated primary antibody (Table 1) overnight at 4°C. After incubation, membranes were washed 3 times for 5 minutes with Tris-buffered saline supplemented with 0.1% Tween-20 (TBS-T). Membranes were then incubated with the indicated secondary antibody for 1 hr at room temperature. After incubation, membranes were washed 3 times for 5 minutes with TBS-T prior to incubation in ECL reagent for 1 minute and imaging using a Chemidoc MP (BioRad). Images acquired were analyzed using Image Lab software (BioRad).

**Table 1.**

| Antibody | Manufacturer | Catalog number |
| --- | --- | --- |
| β-Actin | Santa Cruz Biotechnology | SC-47778 |
| Vinculin | Santa Cruz Biotechnology | SC-73614 |
| Total Rb | Santa Cruz Biotechnology | SC-50 |
| Phospho-Rb (Ser807/811) | Cell Signaling Technology | CS-8516 |
| TUFM | Invitrogen | PA5-27511 |
| ClpP | Cell Signaling Technology | CS-14181 |
| Phospho-AMPK (Thr172) | Cell Signaling Technology | CS-2535 |
| DR5 | ProScience | 2019 |
| Phospho-H2A.X (S139) | Abclonal | AP0687 |
| Total H2A.X | Abclonal | A11463 |
| Phos-CDK1 (Tyr15) | Cell Signaling Technology | CS-2543 |
| Wee1 | Cell Signaling Technology | CS-13084 |
| GDF15 | Santa Cruz Biotechnology | SC-377195 |
| IL-10 | Cell Signaling Technology | CS-12163 |

**Table 2.**
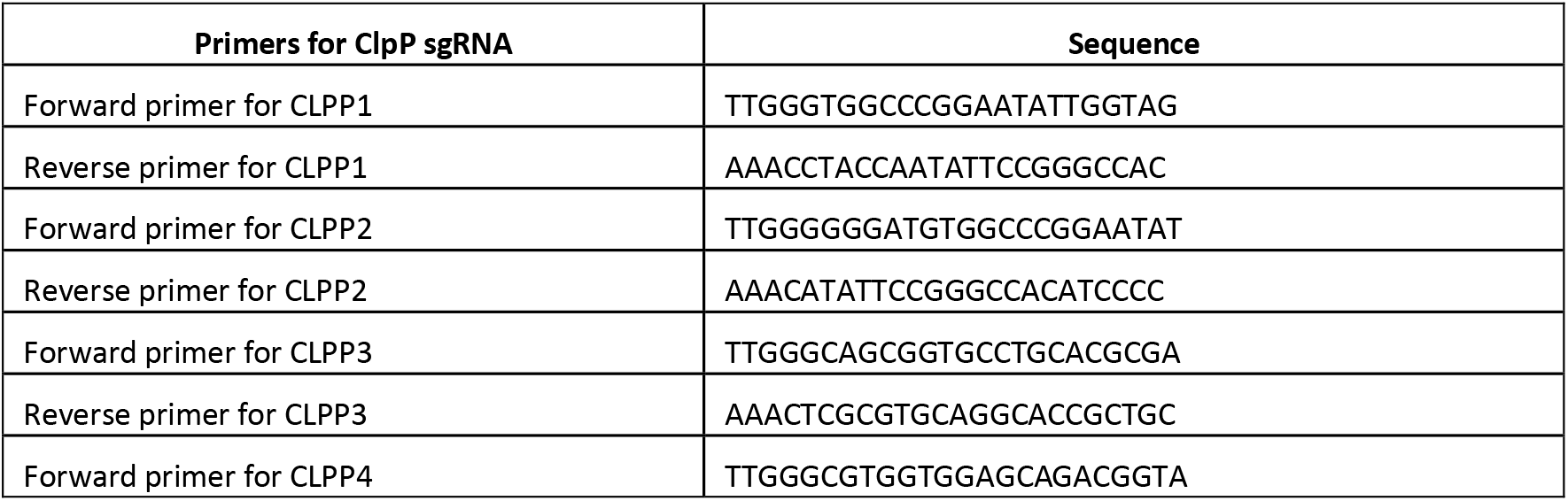

### Generation of ClpP null cells using CRISPRi

To generate sgRNA for CRISPRi, primer pairs for each individual sgRNA were first annealed and later ligated to a digested VDB783 vector (50 ng/μL). Each ligation product was then transformed into DH5α by mixing 3 μL of DNA into 25 μL of competent cells. Cells/DNA mixture was incubated on ice for 30 minutes and heat shocked at 42°C for 45 seconds. Bacteria were centrifuged at maxspeed for 1 minute at room temperature. The resulting bacterial pellet was resuspended in 30 μL of LB, spread on an LB-Amp plate and incubated at 37°C overnight. Colony PCR was performed to check for positive clones of each sgRNA. Single positive clones were grown in 5 mL of LB supplemented with Ampicillin at 37°C overnight. Cultures were miniprepped the next day using a QIAprep Spin Miniprep Kit (Qiagen, U.S.A.) according to the manufacturer’s protocol. Lentivirus were produced in HEK293T cells. Transfection and clonal isolation of the CRISPR null mammalian cells was done as previously described^33^.

### RNA extraction and cDNA synthesis

Total RNA was extracted and purified using RNeasy Mini Kit (Qiagen, U.S.A.) according to the manufacturer’s protocol. cDNA was synthesized from reverse transcription on 1.0 μg total RNA in a 20 μL reaction using High-Capacity cDNA Reverse Transcription Kit (Applied Biosystems, U.S.A.) and T100 thermal cycler (BIO-RAD, U.S.A.), according to the manufacturer’s protocol.

### Quantitative real-time PCR (qRT-PCR)

The cDNA was analyzed by real-time qPCR using iTaq Universal SYBR Green Supermix (BIO-RAD) on an Applied Biosystems 7500 Fast Real-Time PCR System. For each reaction, 1 μL of cDNA was mixed with 12.5 μl of 2 x SYBR mix, 8 μL of water, 1.75 μl of Forward primer and 1.75 μl of Reverse primer. Expression of GAPDH or β-Actin was used to normalize expression of genes of interest. Every biological replicate was analyzed in technical duplicate. Primer targets and sequence are listed in Table 3.

**Table 3.**
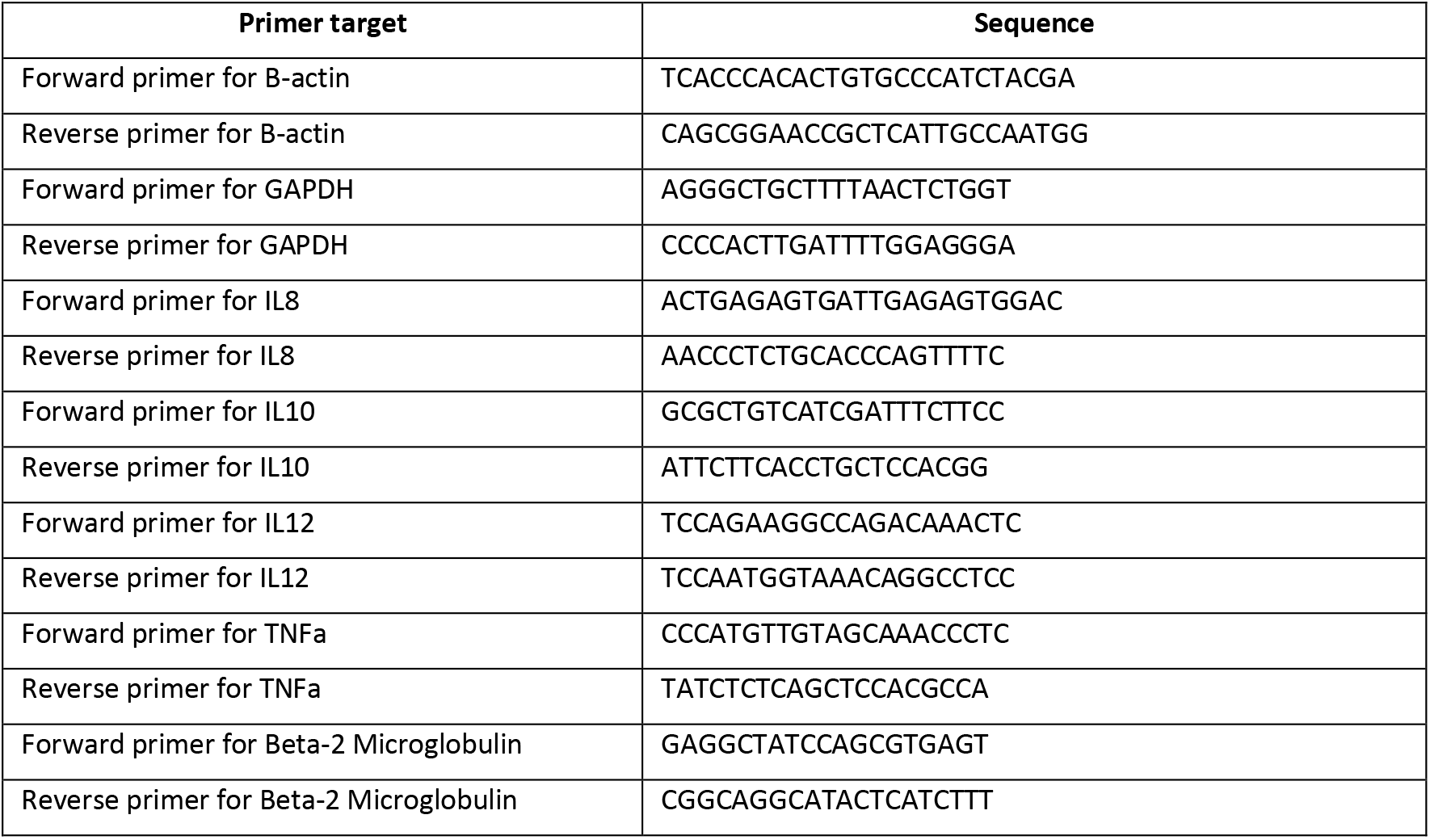

### Caspase Activity Assay

Caspase activity was analyzed using a fluorescent peptide substrate for caspase 3 (Ac-DEVD-AMC) or Caspase 8 (Ac-VETD-AMC). Cells were plated in a 6 well plate (100,000 cells/well) and treated with 0.1% DMSO, 150 nM TR-57, 100 nM staurosporine for 48hrs or 100ng/ml of TRAIL for 6hrs. The samples were harvested by mechanical scraping of the cells into 200 μL of lysis buffer [50 mM HEPES (pH 7.4), 5 mM CHAPS, and 5 mM DTT], lysates were clarified by centrifugation at 10,000xg for 5 minutes at 4°C and protein concentrations were normalized using a Bradford Assay. 100 μg of protein was added to 200 μL of assay buffer [20 mM HEPES (pH 7.4), 0.1% CHAPS, 2 mM EDTA, 5 mM DTT, and 15 μM caspase substrate] in a 96 well plate. The plate was then incubated at room temperature in the dark for 1 hour. The fluorescence intensity from liberated AMC was measured using 360 nm excitation and 460 nm emission filters on a Biorad plate reader. The results were analyzed using GraphPad Prism 9 software.

### Statistical Analysis

Statistical calculations for all the data were performed using GraphPad Prism 9. Data are reported as the mean ± standard error of the mean (S.E.M). S.E.M. was performed on all datasets to determine positive and negative error. Unpaired Two-tailed student t-test or One-way ANOVA was used to make comparisons between groups, and p values below 0.05 at the 95% confidence level were considered to be statistically significant.

## Results

### ClpP activators ONC201 and TR-57 induce senescence in TNBC cell lines

Senescence is phenotypic cell state in which the cell has exited the cell cycle and ceases to proliferate. Increased levels of DNA damage, lack of proliferation, reduced Rb phosphorylation, and increased B-gal activity are all hallmarks of cellular senescence^21,29,30,34,35^. In our previous study, we demonstrated that treatment with ONC201 and the related TR compounds caused growth arrest of the TNBC cell line SUM159 without a reduction in total cell numbers^9^. Thus, we investigated whether these compounds were inducing senescence in these cells. SUM159 cells were treated with 10 μM ONC201 or 150 nM TR-57 for 48 hrs then fixed and stained for β-gal activity using X-gal as described in Material and Methods. Imaging of these cells showed an increase in β-gal positive cells after treatment with ONC201, TR-57, or the CDK4/6 inhibitor known to induce cellular senescence abemaciclib (Fig. 1A) ^36–38^. Manual counting of cells in each treatment group revealed that ONC201 and TR-57 increased the percentage of B-gal positive cells from ~1% in cells treated with vehicle alone to 41% and 37% respectively. The increase in B-gal positive cells observed in response to these compounds was similar to that found with abemaciclib, 45%, which served as a positive control for cellular senescence. (Fig. 1B).

**Fig. 1.**
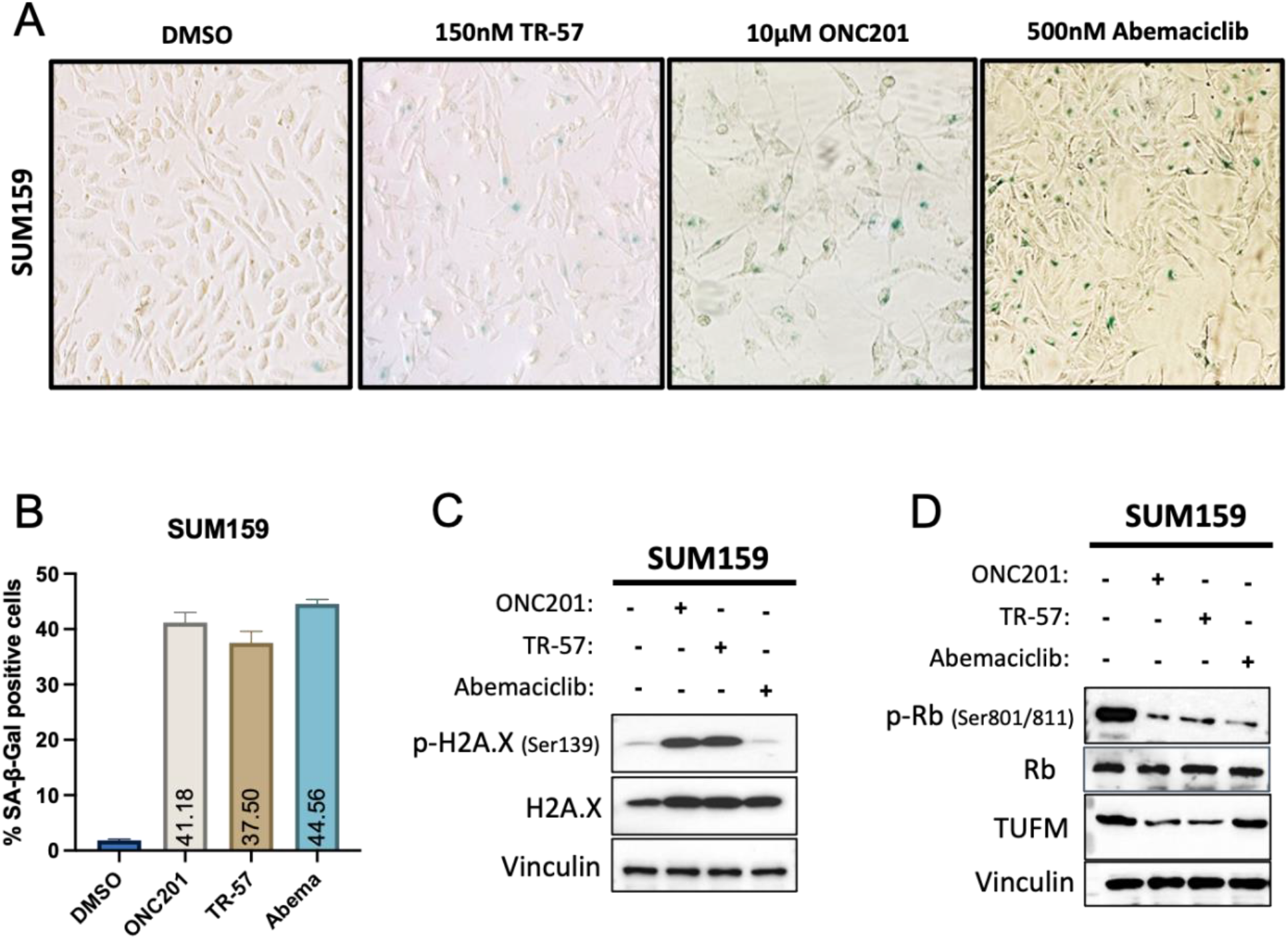
Activation of ClpP induces senescence in TNBC cells. **A**. β-Galactosidase Staining of SUM159 cells 4 days after being treated with ONC201, TR-57, or abemaciclib for 48hrs. **B.** Quantification of images shown in Fig1B. **C.** Immunoblots showing the effect of ONC201, TR-57, and abemaciclib (48hr treatment) on DNA damage marker phospho-H2A.X in SUM159 WT cells. **D.** Immunoblots showing the effect of ONC201, TR-57, and abemaciclib (48hr treatment) on the phosphorylation levels of cell cycle regulator protein Rb in SUM159 WT cells. Data shown in this figure is representative of 3 independent experiments.

We next used immunoblots for phosphorylation marks on H2A.X or Rb to determine whether treatment with ONC201 or TR-57 increased levels of DNA damage or progression through the cell cycle respectively. We observed an increase in H2A.X phosphorylation after ONC201 or TR-57 treatment (Fig. 1C). While CDK4/6 inhibitors induce senescence, they do not increase DNA damage levels^39^. Accordingly, no changes in phosphorylation levels of H2A.X after treatment with abemaciclib were detected (Fig. 1C). ONC201, TR-57, and abemaciclib reduced Rb phosphorylation after 48 hrs (Fig. 1D). Similar results were observed in MDA-MB-231 cells, another TNBC cell line (Supp Fig. 1A). Furthermore, treatment of SUM159 cells with another highly potent ClpP activator^33^, TR107, also caused a strong reduction in Rb phosphorylation (Supp Fig. 2A). As previously described^9,33^, ClpP activation with ONC201 or TR-57 resulted in the loss of the mitochondrial protein TUFM but, as expected, abemaciclib had no effect on TUFM protein levels (Fig. 1D). Together these data indicate that ClpP activators like ONC201 or TR-57 lead to an increase in established senescence markers in these cells.

### Induction of senescence by TR-57 is ClpP dependent

Since ONC201 and TR-57 were equally effective at inducing senescence in both SUM159 and MDA-MB-231 cell lines, we chose to use SUM159 in subsequent experiments. To confirm that TR-57 was inducing senescence through a ClpP-dependent mechanism, we generated SUM159 ClpP null cells using an established dCas9-KRAB system^40,41^. Knockdown of ClpP was verified by immunoblotting. Successful knockdown of ClpP was achieved in ClpP null cells as there was no detectable signal in immunoblots (Fig2A). Knockdown of ClpP did not affect the proliferation of these cells as the doubling time (~21 hours) was determined to be equivalent to the wild type cells. Consistent with the loss of ClpP in ClpP null cells, TUFM protein levels were not reduced upon treatment with TR-57. (Fig. 2A). When compared to wild-type cells, ClpP null cells were largely insensitive to TR-57 in MTS-based viability assays with a >100-fold shift in IC50 (Fig. 2B).

**Fig. 2.**
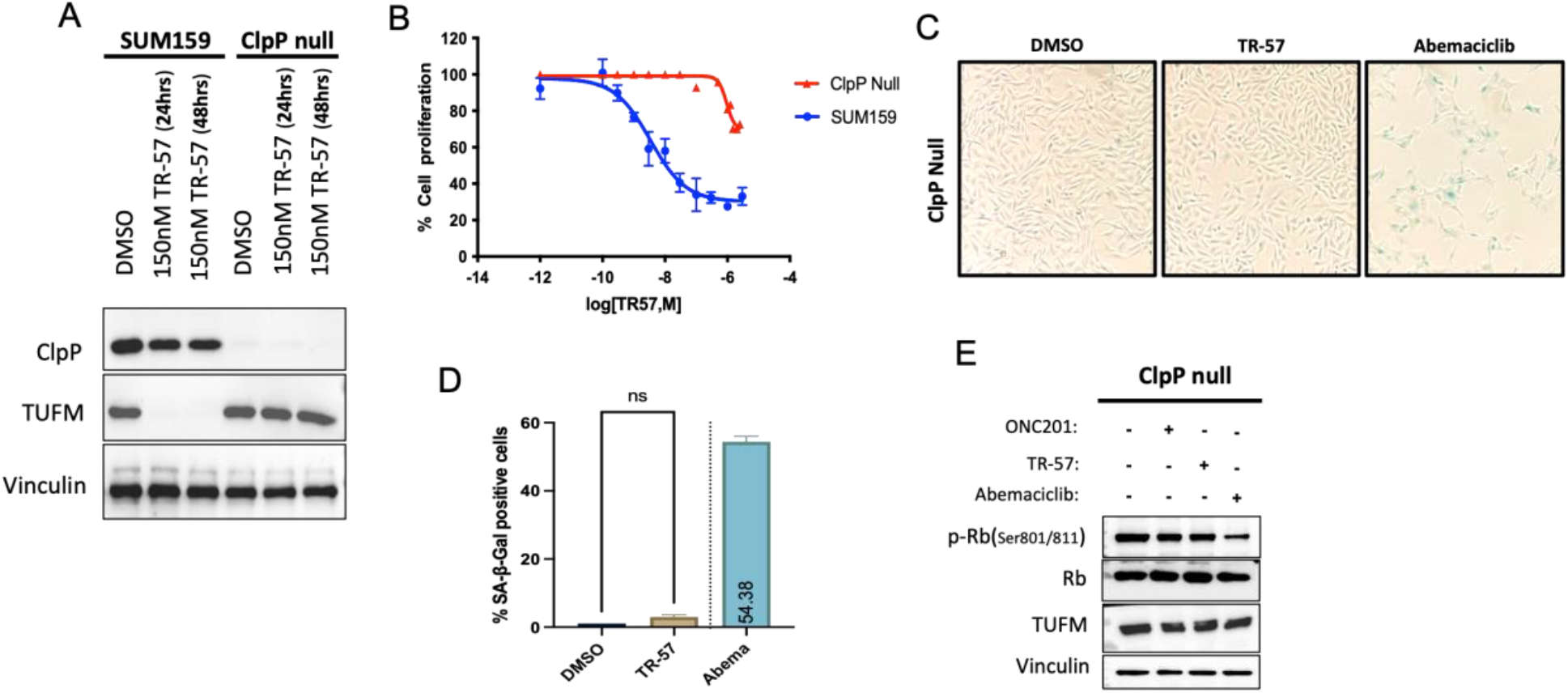
Induction of senescence by TR-57 is ClpP dependent. **A.** Validation of SUM159 ClpP null cell line generated using CRISPRi. Total levels of ClpP protein were assessed in SUM159 WT and ClpP null cells using immunoblots. **B**. Cell viability plots for SUM159 WT and ClpP null cells using MTS assay after 48 hrs treatment with TR-57 (150 nM) **C.** Senescence β-Galactosidase Staining of SUM159 ClpP null cells 4 days after being treated with TR-57 (150 nM) or abemaciclib (500 nM) for 48 hrs. **D.** Quantification of images shown in Fig. 2C. **E.** Immunoblots showing the effect of ONC201 (10 μM), TR-57 (150 nM), and abemaciclib (500 nM) 48 hrs treatments on the phosphorylation levels of cell cycle regulator protein Rb in SUM159 ClpP null cells. Data shown in this figure is representative of 3 independent experiments.

Next, we investigated whether TR-57 induced senescence in ClpP null cells by evaluating the parameters described above. While abemaciclib induced senescence in the SUM159 ClpP null cells, no induction of senescence was observed after incubation with TR-57 as shown by the lack of β-gal positive cells (Fig. 2C). Notably, percent of β-gal positive ClpP null cells after abemaciclib (54%) (Fig. 2D), was similar to that observed with wild type SUM159 cells (44%) (Fig. 1B). Moreover, TR-57 did not reduce Rb phosphorylation, whereas abemaciclib strongly reduced Rb phosphorylation in ClpP null cells (Fig. 2E). These data demonstrate that induction of senescence by TR-57 in TNBC cells is ClpP dependent and differs from mechanisms driving senescence upon treatment with abemaciclib.

### TR-57 increases immune markers associated with senescence

IL-8 has been shown to regulate inflammatory responses and plays an important role as a leukocyte activator when secreted by senescent cells^25,26^. Thus, IL-8 is considered a key component of SASP. Induction of MiDAS has been reported to increase the levels of anti-inflammatory and pro-inflammatory cytokines IL-10 and TNFa, respectively^27,42^. To test whether any of these senescence associated cytokines were upregulated upon treatment with TR-57, SUM159 cells were treated with TR-57 as described above and IL-8, IL-10, IL-12, and TNFa mRNA levels were measured by qRT-PCR as described in Material and Methods. TR-57 induced a 3-fold increase in IL-8, a 7-fold increase in IL-12, and a 28-fold increase in IL-10 (Fig. 3A). Additionally, TR-57 caused a 2-fold decrease in TNFa mRNA.

**Fig. 3.**
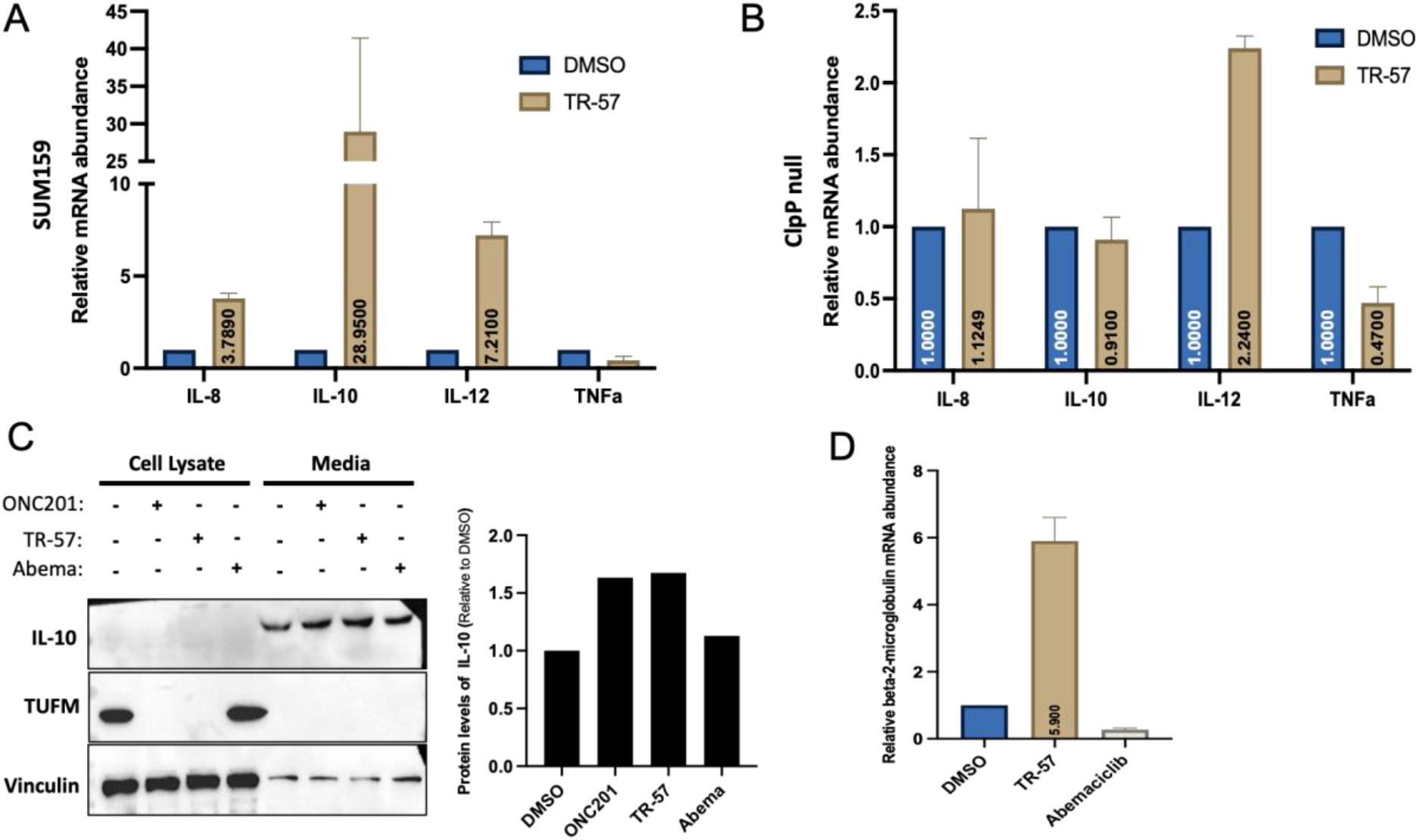
ClpP activation increases immune markers associated with senescence. **A.** Gene expression of immune markers IL-8, IL-10, IL-12 and TNFa in SUM159 WT cells after being treated with TR-57 (150 nM) for 48 hrs. **B**. Gene expression of immune markers IL-8, IL-10, IL-12 and TNFa in SUM159 ClpP null cells after being treated with TR-57 (150 nM) for 48 hrs. **C.** Immunoblots showing the effect of ONC201 (10 μM), TR-57 (150 nM), and abemaciclib (500 nM) 48 hr treatments on IL-10 protein levels in SUM159 WT cells **D.** Gene expression of immune marker B2M in SUM159 WT cells after being treated with TR-57 (150 nM) or abemaciclib (500 nM) for 48 hrs.

To confirm that changes in cytokine expression were ClpP dependent, the experiment was repeated in ClpP null cells. TR-57 had no effect on IL-8 or IL-10 mRNA levels in the ClpP null cells (Fig. 3B). A 2-fold increase in IL-12 mRNA expression was observed in the ClpP null cell after TR-57 treatment when compared to the DMSO control, suggesting that ClpP loss only partially suppressed IL-12 upregulation. Also, TR-57 treatment led to a ~2-fold decrease in TNFa mRNA. These experiments suggested that changes by TR-57 in TNFa and IL-12 mRNA where not completely ClpP dependent while the increase in IL-8 and IL-10 mRNA was ClpP dependent. These ClpP dependent events are consistent with the development of senescence as determined by β-gal and other cellular markers (Fig. 2C & 2E). In order to validate our qRT-PCR data, we measured the amount of IL-10 protein in SUM159 cells and in tissue culture supernatant by immunoblot. While no significant levels of IL-10 were detected in cell lysates an increase in IL-10 protein was detected in the media of cells that were treated for 48 hrs with ONC201, TR-57, or abemaciclib, when compared to DMSO (Fig. 3C). These results demonstrate that TR-57 is inducing a SASP response in SUM159.

AMPK activation is an established marker of MiDAS^27^ in addition to upregulation of IL-10 shown above. We next evaluated if TR-57 treatment increased AMPK phosphorylation in SUM159 cells. As determined by immunoblotting for phospho-AMPK, TR-57 increased phospho-AMPK levels after 48 hrs (Supp Fig. 3A). We also evaluated the mRNA levels of Beta-2-microglobulin (B2M) after TR-57 treatment, as senescent cells have been shown to trigger anti-tumor immunity through upregulation of B2M and other class I major histocompatibility complex members^37^. Quantification of B2M mRNA levels demonstrated that TR-57 induced a ~6-fold increase when compared to the DMSO control (Fig. 3D). Lastly, since growth differentiation factor 15 (GDF15) is a marker associated with aging and senescence in multiple cell models^43^, we evaluated the effects of ClpP agonists on GDF15 protein levels. As determined by immunoblotting, both ONC201 and TR-57 increased GDF15 levels at 24 and 48 hrs (Supp Fig. 3B/C). Altogether, our data suggest that TR-57 is inducing events associated with MiDAS in SUM159 cells.

### TR-57 alters the expression of antiapoptotic/proapoptotic proteins

Previous reports have shown that senescent cells alter pro-survival responses through decreased expression of pro apoptotic proteins^44,45^. Therefore, we examined if induction of senescence by TR-57 affected the levels of apoptosis-related proteins in SUM159 cells using a human apoptosis array kit (R&D Systems, Inc., USA). Treatment with TR-57 led to a decrease in XIAP, Survivin, and SMAC/DIABLO (Fig. 4A). Interestingly, TRAIL receptor 1 (DR4), and TRAIL receptor 2 (DR5) were also downregulated when compared to the DMSO treated sample (Fig. 4B). The reduction in DR5 was further validated by immunoblotting. As shown in figure 4C, treatment with TR-57 or abemaciclib led to a significant decrease of DR5 protein levels. Next, we tested whether TR-57 treatment had an effect on DR5 mRNA levels using qRT-PCR. Quantification of DR5 mRNA levels demonstrated that TR-57 induced a ~1.5-fold decrease compared to the DMSO control (Figure 4D). These findings suggest that TR-57 suppresses expression of proapoptotic protein levels and reduces gene expression of DR5.

**Fig. 4.**
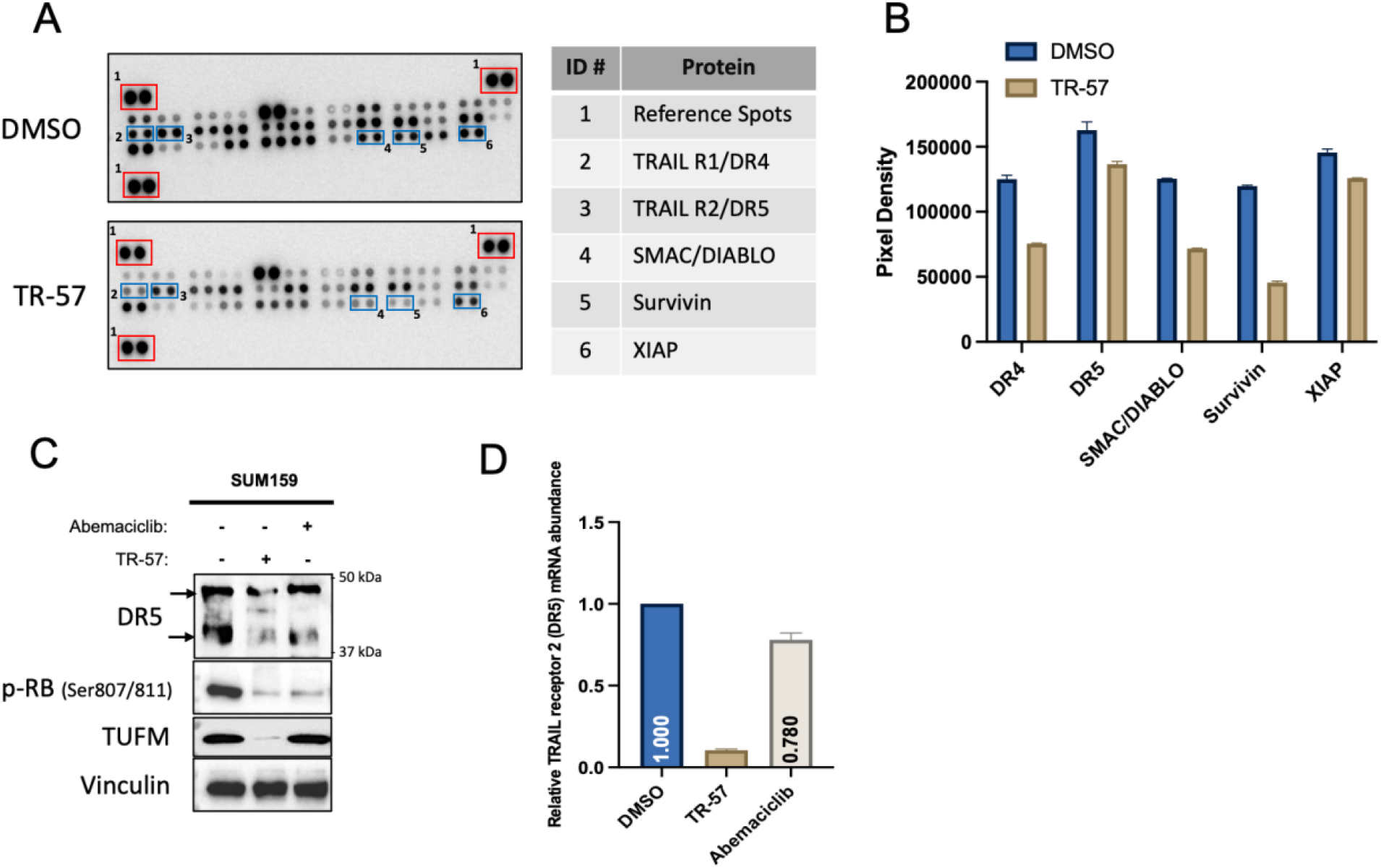
Induction of senescence leads to downregulation of apoptotic proteins. **A.** Detection of apoptosis related proteins levels in SUM159 cells after TR-57 (150 nM) treatment for 48 hrs using a human apoptosis array **B**. Quantification of DR4/5 array blot data shown in Fig. 4A **C**. Immunoblots showing the effect of TR-57 (150 nM) or abemaciclib (500 nM) treatments for 48 hrs on DR5 protein levels in SUM159 WT cells. **D.** Gene expression of DR5 in SUM159 WT cells after 48 hrs treatment of TR-57 (150 nM) or abemaciclib (500 nM).

### Induction of senescence sensitizes TNBC cells to TRAIL induced apoptosis

Downregulation of proapoptotic proteins like SMAC/DIABLO or DR5 suggested that cells exposed to ClpP activators may be more resistant to apoptosis induction. Addition of TR-57 alone did not increase caspase-3 activity, whereas treatment with staurosporine (STS) or TRAIL led to a ~6-fold and ~5-fold increase in caspase-3 activity, respectively (Fig. 5A). Notably, treatment with TR-57 followed by STS prevented the increase in caspase-3 activity previously observed with STS alone. (Supp Fig. 4A)

**Fig. 5.**
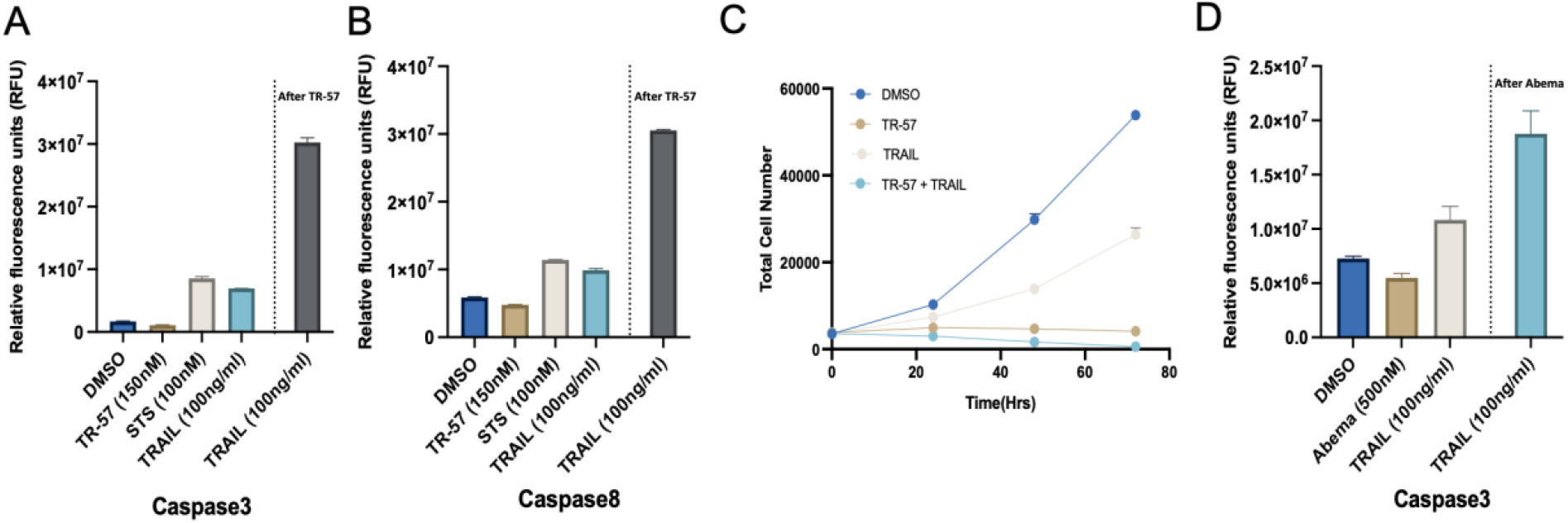
Senescent cells show increased sensitivity to TRAIL-induced apoptosis. **A.** Measurement of caspase 3 activity in SUM159. Cells were treated with TR-57 (150 nM) for 48 hrs, STS (100 nM) for 24 hrs, TRAIL (100 ng/ml) for 6 hrs, or TR-57 (150 nM) for 48 hrs in combination with TRAIL (100ng/ml) for 6 hrs **B**. Measurement of caspase 8 activity in SUM159. Cells were treated as described in Fig. 5A **C.** Total cell count assay of SUM159 cells. Cells were treated with TR-57 (150 nM) for 48 hrs, TRAIL (100 ng/ml) for 6 hrs, or TR-57 (150 nM) for 48 hrs in combination with TRAIL (100ng/ml) for 6 hrs and imaged following Hoechst stain addition after 72 hours. **D.** Measurement of caspase 3 activity in SUM159 after treatment with abemaciclib alone or in combination with TRAIL. Caspase activity for Fig. 5 A, B, and C was measured using a fluorescent peptide substrate assay as described in methods. Data shown in this figure is representative of 3 independent experiments.

Since ONC201 was previously reported to induce TRAIL expression and caspase-dependent apoptotic cell death through DR5 activation^37–39^, we next compared the effects of TRAIL addition after TR-57 exposure. Interestingly, treatment with TR-57 followed by TRAIL led to a ~26-fold increase in caspase-3 activity, when compared to TR-57 alone (Fig. 5A). Similar results were obtained using a caspase-8 assay (Fig. 5B). Consistent with drastically increased caspase activity, cell death was confirmed by measuring total cell number after TRAIL treatment alone or in combination with TR-57 (Fig. 5C). Treatment with TRAIL alone was sufficient to inhibit cell proliferation by about 50% whereas no proliferation was observed after 72 hours of TR-57. By contrast, the combination of TR-57 followed by TRAIL resulted in a reduction of cell number from 4000 on day 0 to only 582 following 72 hours of treatment suggesting cell death and consistent with enhanced caspase activity. Similar results were observed in another TNBC cell line, MDA-MB-231(Supp Fig. 5A).

Lastly, we compared whether the increased sensitivity to TRAIL-induced apoptosis was affected by an alternative senescence inducer. While treatment with abemaciclib alone did not increase caspase-3 activity, the combinatorial treatment of abemaciclib followed by TRAIL resulted in an ~ 1.5-fold increase in caspase-3 activity (Fig. 5D). In summary, these results confirmed that senescence induction by TR-57 or abemaciclib leads to an increase in sensitivity of TNBC cells to TRAIL induced apoptosis.

### Downregulation of Wee1 partially mediates TNBC cells sensitization to TRAIL induced apoptosis

Previous studies have shown that loss or inhibition of Wee1 leads to TRAIL sensitization in breast cancer cells^48,49^. Therefore, to expand upon our recent results, we evaluated the effects of TR-57 on Wee1 protein levels. Treatment with TR-57 or abemaciclib reduced total Wee1 protein levels in a ClpP dependent manner (Fig. 6A). We next tested whether adavosertib, a Wee1 inhibitor, also resulted in an increase in TRAIL sensitivity. While treatment with adavosertib alone did not increase caspase-3 activity, combinatorial treatment of adavosertib followed by TRAIL incubation, led to a ~2-fold increase in caspase-3 activity when compared to DMSO and a ~0.7-fold increase when compared to TRAIL alone (Fig. 6B). Successful inhibition of Wee1 after adavosertib treatment was verified by the decrease of CDK1 phosphorylation, an established Wee1 substrate (Supp Fig. 6A). These data suggest that reduction of Wee1 in response to senescence induction may contribute to the enhanced effects of TRAIL in these cells.

**Fig. 6.**
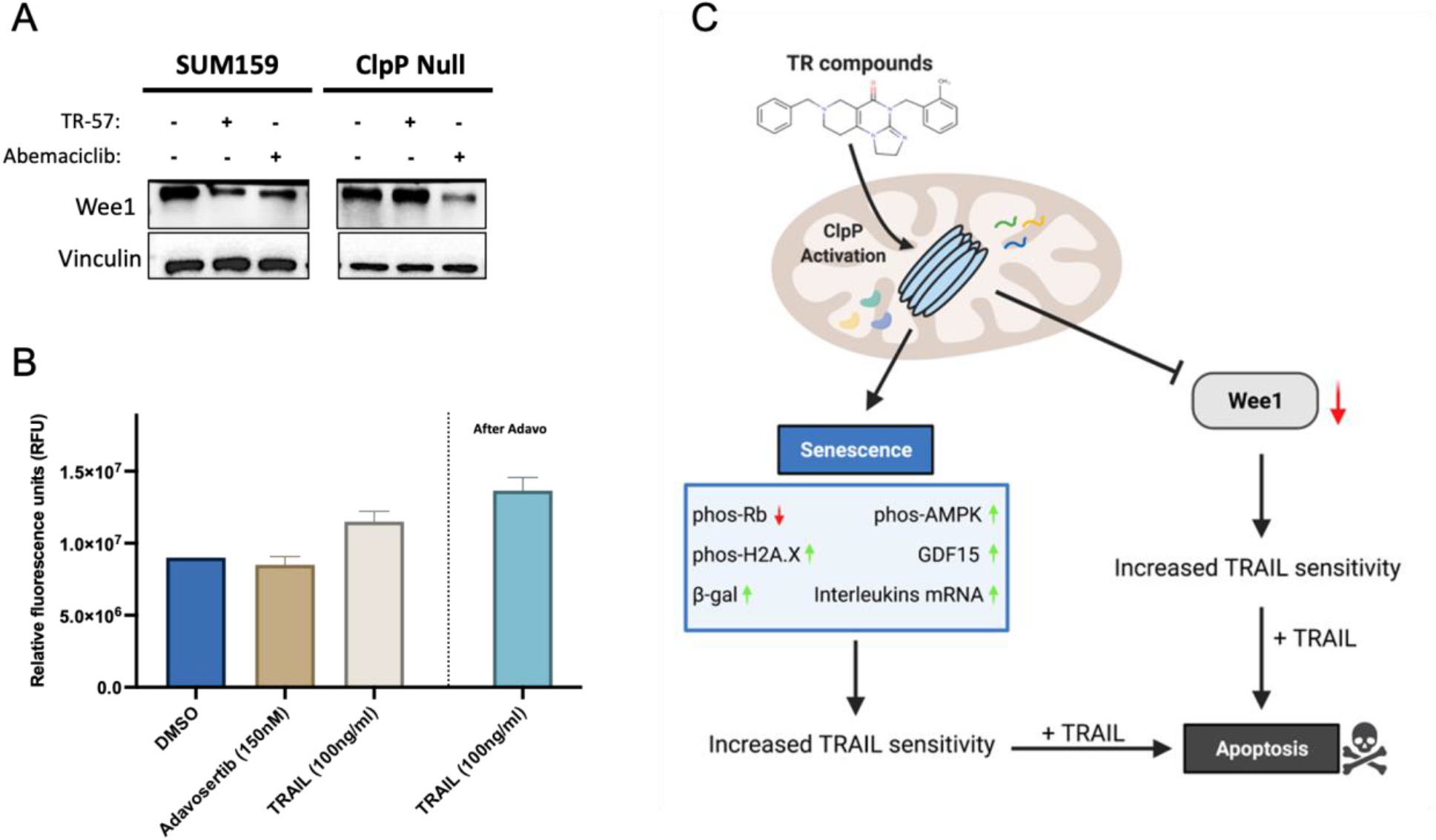
Downregulation of Wee1 partially mediates TNBC cells sensitization to TRAIL-induced apoptosis. **A.** Immunoblots showing the effect of TR-57 (150 nM) or abemaciclib (500 nM) treatments for 48 hrs on Wee1 levels in SUM159 WT and SUM159 ClpP null cells. **B**. Measurement of caspase 3 activity in SUM159. Cells were treated with Adavosertib for 24 hrs, STS (100 nM) for 24 hrs, TRAIL (100 ng/ml) for 6 hrs, or Adavosertib for 48 hrs in combination with TRAIL (100 ng/ml) for 6 hrs. **C.** Proposed model on how ClpP agonists lead to senescence and TRAIL sensitivity. Data shown in this figure is representative of 3 independent experiments.

## Discussion

The discovery of ONC201 and related analogs has stimulated considerable interest in small molecule ClpP activators as novel anti-cancer agents. While the effects of ONC201 were initially attributed to effects on TRAIL and later dopamine receptors^5,12,46,47^, inconsistencies in the literature argued against these as common mechanisms of action. Importantly, differential effects on apoptosis, kinase signaling, TRAIL induction and dopamine receptor signaling were observed across multiple cancer models further suggesting cell type dependent responses. In contrast to other cancer models, TNBC cells showed little or no apoptosis in response to ONC201 and related analogs^9,13^. In fact, these studies suggested a cytostatic response that was not dependent on TRAIL induction or expression of TRAIL receptors^13^. The results of our studies not only confirm that activation of ClpP has a cytostatic effect on cell growth but demonstrate a corresponding increase in senescence markers in TNBC cells.

In this report we also compared and contrasted the effects of abemaciclib, an established inducer of senescence in breast cancer models^37^. abemaciclib and other CDK4/6 inhibitors have been shown to effectively increase senescence in other TNBC models. Our studies demonstrate that TR-57, a highly potent and selective activator of the mitochondrial protease ClpP, induced established markers of senescence including increases in DNA damage and similar SASP profiles. Induction of senescence by TR-57 was equivalent to that observed with CDK4/6 inhibition but was completely dependent on ClpP whereas abemaciclib was not dependent on ClpP. This contrasted with the effects of the chemotherapeutic doxorubicin, which induced senescence in a manner partially dependent on ClpP (data not shown). In addition to evoking SASP, TR-57 led to an increase in gene expression of immune marker B2M. This data is consistent with findings that ClpP activators can promote immune recruitment *in vivo*^50^.

Increased sensitivity to TRAIL may be of pivotal importance to the effects of ClpP activators on the balance between senescence and apoptosis. Recent studies have shown that treatment with ClpP activators sensitizes endometrial, pancreatic ductal adenocarcinoma and other cancer cells to TRAIL induced apoptosis^51–53^. Our study shows that increased TRAIL sensitivity after ClpP activators is also observed in TNBC cells. However, most of the recent studies attributed such change in sensitivity to an increase in DR5 receptors. In this regard our study differs in that we observed reduced DR5 protein and mRNA levels in response to TR-57 treatment while increasing TRAIL mediated apoptotic response. Therefore, our findings argue that upregulation of DR5 is not necessary in TNBC models and that there is a potent TRAIL response even at reduced levels of DR5 expression induced by TR-57 treatment. Lastly, our study demonstrates that senescent TNBC cells had lower levels of Wee1, that correlated with an increase in senescence. While treatment with the Wee1 inhibitor adavosertib led to an increase in TRAIL sensitivity, the changes observed were modest when compared to the increase observed after TR-57. Thus, our data suggest that Wee1 inhibition may be partially responsible for the shift in TRAIL sensitivity but that additional mechanisms may contribute to the potent modulation of TRAIL sensitivity by ClpP activators.

In conclusion, our study shows that ClpP activators induce senescence in TNBC cell lines in a ClpP dependent manner. Our findings also highlight that combining ClpP activators like TR-57 to induce senescence followed with TRAIL treatment provides an effective approach to arrest TNBC cells growth and eliminate malignant senescent cells *in vitro*.

## Acknowledgements

This project is supported by grants from National Institutes of Health to LMG (5R01GM138520-02). This research was supported in part by the Intramural Research Program of the National Cancer Institute, Center of Cancer Research (ZIA SC 007263 to SL).

