## Supplementary figures for "SMALL MOLECULE CLPP AGONISTS INDUCE SENESCENCE AND ALTER TRAIL-MEDIATED APOPTOTIC RESPONSE OF TRIPLE-NEGATIVE BREAST CANCER CELLS"

### Supplemental Figures

A

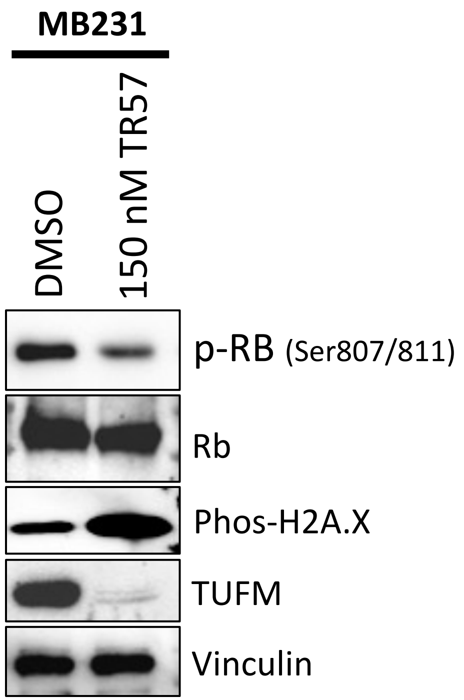


**Sup Fig. 1 Activation of ClpP modulates senescence markers in MDA-MB-231 cells**

**A.** Immunoblots showing the effect of TR57 (150 nM) for 48hrs on DNA damage marker phospho-H2A.X and on the phosphorylation levels of cell cycle regulator protein Rb in MDA-MB-231 WT cells. Data shown in this figure is representative of 2 independent experiments.

A

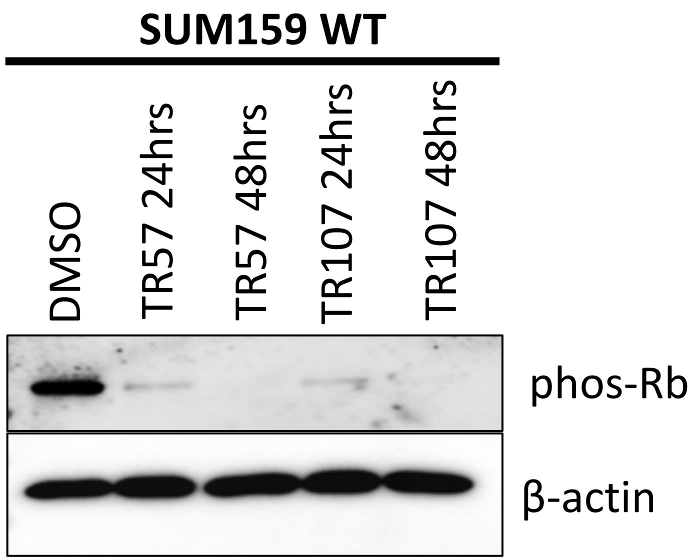


**Sup Fig. 2 Treatment with TR107 leads to loss of Rb phosphorylation**

**A.** Immunoblots showing the effect of TR57 (150 nM) or TR107 on the phosphorylation levels of cell cycle regulator protein Rb in SUM159 WT cells. Data shown in this figure is representative of 2 independent experiments.

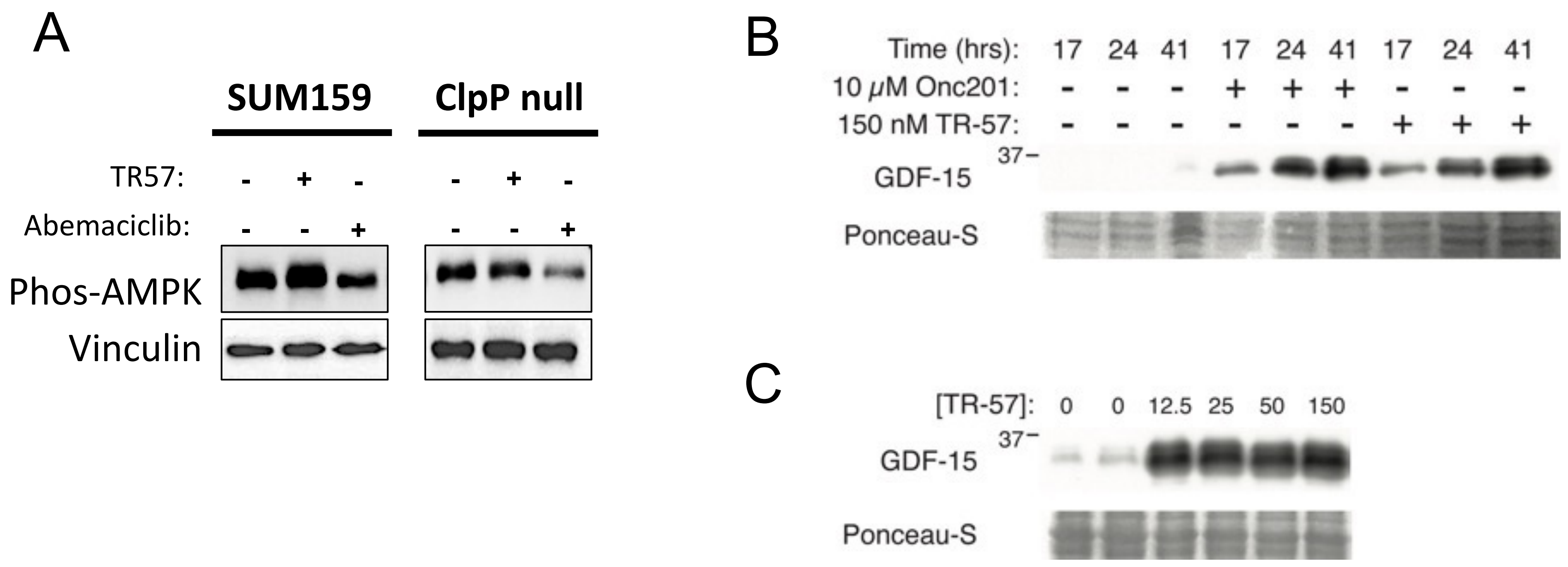


**Sup Fig. 3 Activation of ClpP leads to an increase in AMPK phosphorylation and Total GDF15 levels in SUM159 cells**

**A.** Immunoblots showing the effect of TR57 (150 nM) or Abemaciclib (500 nM) on the phosphorylation levels of AMPK in SUM159 WT and SUM159 ClpP null cells. **B.** SUM159 cells were treated with vehicle (DMSO at 0.1%), ONC201, or TR-57 for the indicated times and immunoblotted for GDF-15. Ponceau-S staining indicates similar protein loading. **C.** SUM159 cells were treated with different concentrations of TR-57 for 24 hours and immunoblotted as described in B. Data shown in this figure is representative of 2 independent experiments.

A

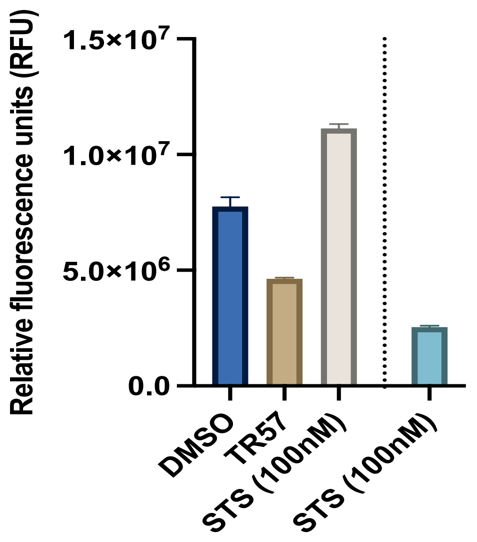


**Sup Fig. 4 Senescent SUM159 cells show decreased sensitivity to STS caspase 3 activation**

**A.** Measurement of caspase 3 activity in SUM159. Cells were treated with TR-57 (150 nM) for 48 hrs, STS (100 nM) for 24 hrs, or TR-57 (150 nM) for 48 hrs in combination with STS (100 nM). Data shown in this figure is representative of 2 independent experiments.

A

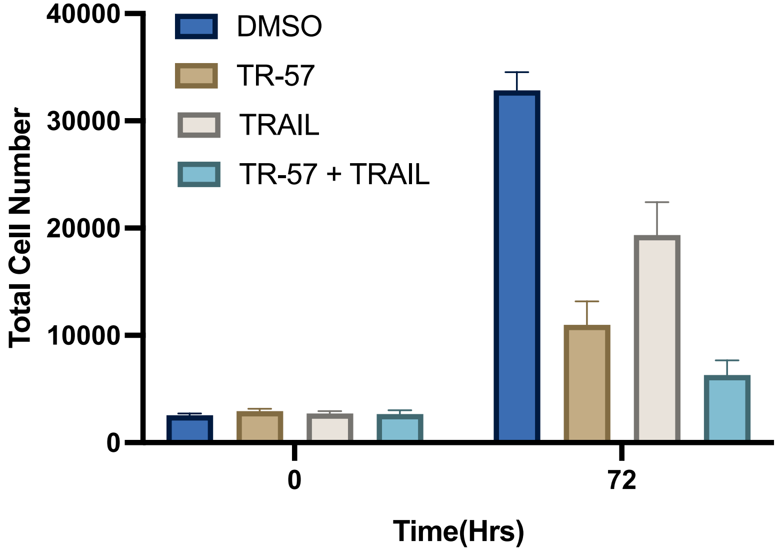


**Sup Fig. 5 MDA-MB-231 cells show increased sensitivity to TRAIL induced apoptosis**

**A.** Total cell count assay of MDA-MB-231 cells. Cells were treated with TR-57 (150 nM) for 48 hrs, TRAIL (100 ng/ml) for 6 hrs, or TR-57 (150 nM) for 48 hrs in combination with TRAIL (100ng/ml) for 6 hrs and imaged following Hoechst stain addition after 72 hours. Data shown in this figure is representative of 2 independent experiments.

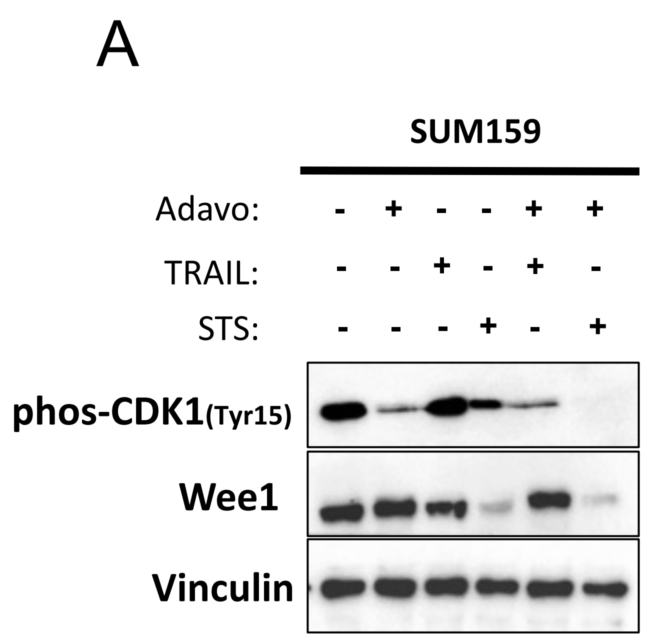


**Sup Fig. 6 Validation of Wee1 inhibition by Adavosertib in SUM159 cells**
**A.** Immunoblots showing the effect of Adavoserib, TRAIL, or STS on the phosphorylation levels of Wee1 substrate CDK1 in SUM159 WT cells. Data shown in this figure is representative of 2 independent experiments.
